# Reconstruction of fish allergenicity from the content and structural traits of the component β-parvalbumin isoforms

**DOI:** 10.1101/813659

**Authors:** Raquel Pérez-Tavarez, Mónica Carrera, María Pedrosa, Santiago Quirce, Rosa Rodriguez-Perez, María Gasset

## Abstract

Most fish-allergic patients have anti-β-parvalbumin (β-PV) immunoglobulin E (IgE), which cross-reacts among fish species with variable clinical effects. Although the β-PV load is considered a determinant for allergenicity, fish species express distinct β-PV isoforms with unknown pathogenic contributions. To identify the role various parameters play in allergenicity, we have taken *Gadus morhua* and *Scomber japonicus* models, determined their β-PV isoform composition and analyzed the interaction of the IgE from fish-allergic patient sera with these different conformations. We found that each fish species contains a major and a minor isoform, with the total PV content four times higher in *Gadus morhua* than in *Scomber japonicus*. The isoforms showing the best IgE recognition displayed protease-sensitive globular folds, and if forming amyloids, they were not immunoreactive. Of the isoforms displaying stable globular folds, one was not recognized by IgE under any of the conditions, and the other formed highly immunoreactive amyloids. The results showed that *Gadus morhua* muscles are equipped with an isoform combination and content that ensures the IgE recognition of all PV folds, whereas the allergenic load of *Scomber japonicus* is under the control of proteolysis. We conclude that the consideration of isoform properties and content may improve the explanation of fish species allergenicity differences.

## INTRODUCTION

Fish is one of the eight major foods causing type I food allergy, an immunoglobulin E (IgE)-mediated hypersensitivity disease resulting from the breakdown of immune tolerance^**1,2,3,4**^. Changes in fishery policies, aquaculture, processing and eating habits have modified the landscape of fish consumption, increasing the worldwide fish allergy prevalence^**3,4**^. The major allergen of fish has been identified as β-parvalbumin (β-PV), a Ca^2+^-binding acidic protein that is abundant in muscle^**3,4,5,6,7**^. Most allergic patients react to more than one fish species due to the cross-reactivity of their β-PV chains, but despite the presence of anti-β-PV, IgE often tolerates a variety of species^**8,9,10,11,12**^. The tolerance of cartilaginous fish results from the absence of cross-reactivity of the basic α-lineage of the expressed PV^**12**^. In contrast, differences in the allergenic potency of bony fish species have been related to both the content and the pattern of the expressed β-PV isoforms^**13,14,15**^. However, the molecular signatures of and the relation between these features are still unknown.

Fish β-PVs are made up of a single polypeptide chain of approximately 109 amino acids that contains three EF-hand motifs (AB, CD and EF), of which only two (the CD and EF) have functional cation-binding acid loops^**3,4,16,17**^. These chains display a complex pattern of IgE linear epitopes that involve intermotif spacers (residues 33-44 joining the AB and CD motifs, and 65-74 linking the CD and EF motifs), acid loops (residues 49-64 and 88-94 on the CD and EF motifs, respectively) and a salmonid-specific region (residues 1-18)^**18,19,20,21,22,23,24,25**^. Of these, the segment containing residues 33-44 has been described as highly immunoreactive in β-PV from several fish species^**3,19,20,24,25**^. Ca^2+^ binding stabilizes the proteins in a helical globular fold, as shown by the 3D structures of cod (Gad m 1) and mackerel (Sco j 1) major β-PVs, among others ^**3,26,27**^. Impairment of this fold in the carp β-PV Cyp c 1 by mutation of both acid loops reduces the IgE binding by 95%, supporting the existence of conformational epitopes^**28**^. However, cation removal by either gastric pH or chelates at neutral pH triggers Gad m 1 aggregation into amyloid assemblies^**25,29**^. Gad m 1 amyloids are formed by the polymerization of the intermotif linker regions and cause the amplification of IgE binding^**25,29,30**^. This sequence and concentration-dependent conformational transition are not unique to this β-PV, having been described for a large number of proteins, some of which, such as β-lactoglobulin (Bos d 5), ovalbumin (Gal d 2) and κ-casein (Bos d 8), are food allergens^**31, 32**^.

Commonly consumed fish are restricted to a few orders, including *Clupeiformes* (herring, sardine, and anchovy), *Cypriniformes* (carp), *Gadiformes* (cod, pollock, and hake), *Perciformes* (perch, snapper, tuna, mackerel, and tilapia), *Pleuronectiformes* (sole and whiff), and *Salmoniformes* (salmon, trout, and whitefish)^**33,34**^. Fish species from these orders differ in the total content of β-PV, the pattern of the expressed isoform and the tolerance in allergic patients^**11,12,35,36,37,38,39**^. Of them, cod *(Gadus morhua*), mackerel (*Scomber japonicus*) and tuna (*Thunnus albacares)* are examples of fish species with high (>2.5 mg/g muscle), medium (0.3-0.7 mg/g muscle) and low (<0.5 mg/g muscle) β-PV content, respectively ^**12,13,14,34,40,41**^. In addition, for a given species, the β-PV concentration varies with fish part, and it is higher in dorsal regions than in ventral regions and in the rostral than in the caudal regions^**40**^. Examination of the UniProtKB sequence base shows that the *Gadus morhua* β-PV family is composed of gmPV1 (A5I874, Gad m 1.0202), gmPV2 (A5I873, Gad m 1.0102) and single residue variants of each of the chains (Q90YK9 and Q90YL0). Of these isoforms, gmPV1 appears to govern the IgE-binding properties of the population isolated from cod muscle^**37**^. For *Scomber japonicus* displaying heat-sensitive allergenicity, two chains have been described so far: conventional sjPV1 (Sco j 1, Q3C2C3) and sjPV2 (Sco j 1, Q9I591), which have been detected at the transcriptional level^**42**^. For *Thunnus albacares*, UniProtKB uniquely reveals the C6GKU3 sequence (Thu a 1). Of these three species, mackerel and tuna are more highly tolerated than cod^**11,12**^.

To simultaneously evaluate the role that content and β-PV isoforms play in the PV allergenic potency of fish species, we chose *Gadus morhua* and *Scomber japonicus* models given their difference in allergen load and the availability of two protein sequences and analyzed the interaction of the IgE of fish-allergic patient sera with the denatured, globular and fibrillary folded states of the β-PVs. The results obtained provide novel variables that can be included in predictions of clinically relevant cross-reactivity from diagnostic tests.

## RESULTS

### Sequence features of *Gadus morhua* and *Scomber japonicus* β-PV isoforms

The sequences of the β-PV isoforms from *Gadus morhua* (gmPV1, gmPV2) and *Scomber japonicus* (sjPV1 and sjPV2) are shown in **Fig. 1a**, together with their pairwise identity patterns (**Fig. 1b**) and the location of relevant immunological (**Fig. 1c**) and structural regions (**Fig. 1d**). Analysis with BioEdit alignment tools showed 56% global identity and 86% global homology among the isoforms. The pairwise identity of proteins varied from 70.6% in *Gadus morhua* to 81.6% in *Scomber japonicus*. Sequence differences were mainly found at the N-terminal half of the chains, overlapping with the predicted PARV19^**36**^ epitope, the most common IgE immunoreactive region (IgE-I)^**3,4,25,43**^, the N-terminal EF-hand (AB region) and the amyloid forming segments of gmPV1^**25,30**^. Despite the sequence differences, the AmyloPred2 algorithm revealed several regions with aggregation potential in all chains (**Fig. 1d**). The chains differed in the number of Cys residues. All chains shared C19, three of the chains (gmPV2, sjPV1 and sjPV2) also contained C34, and gmPV2 also had C12, which suggested differences in their capacity to form disulfide-bonded species and aggregated polymorphs ^**44**^. It must also be noted that K29, which is only present in sjPV1, was suggested to be part of an IgE conformational epitope in this protein^**43**^. All chains were produced as recombinant proteins and purified as Ca^2+^-bound forms using established protocols^**25,45**^.

**Figure 1.**
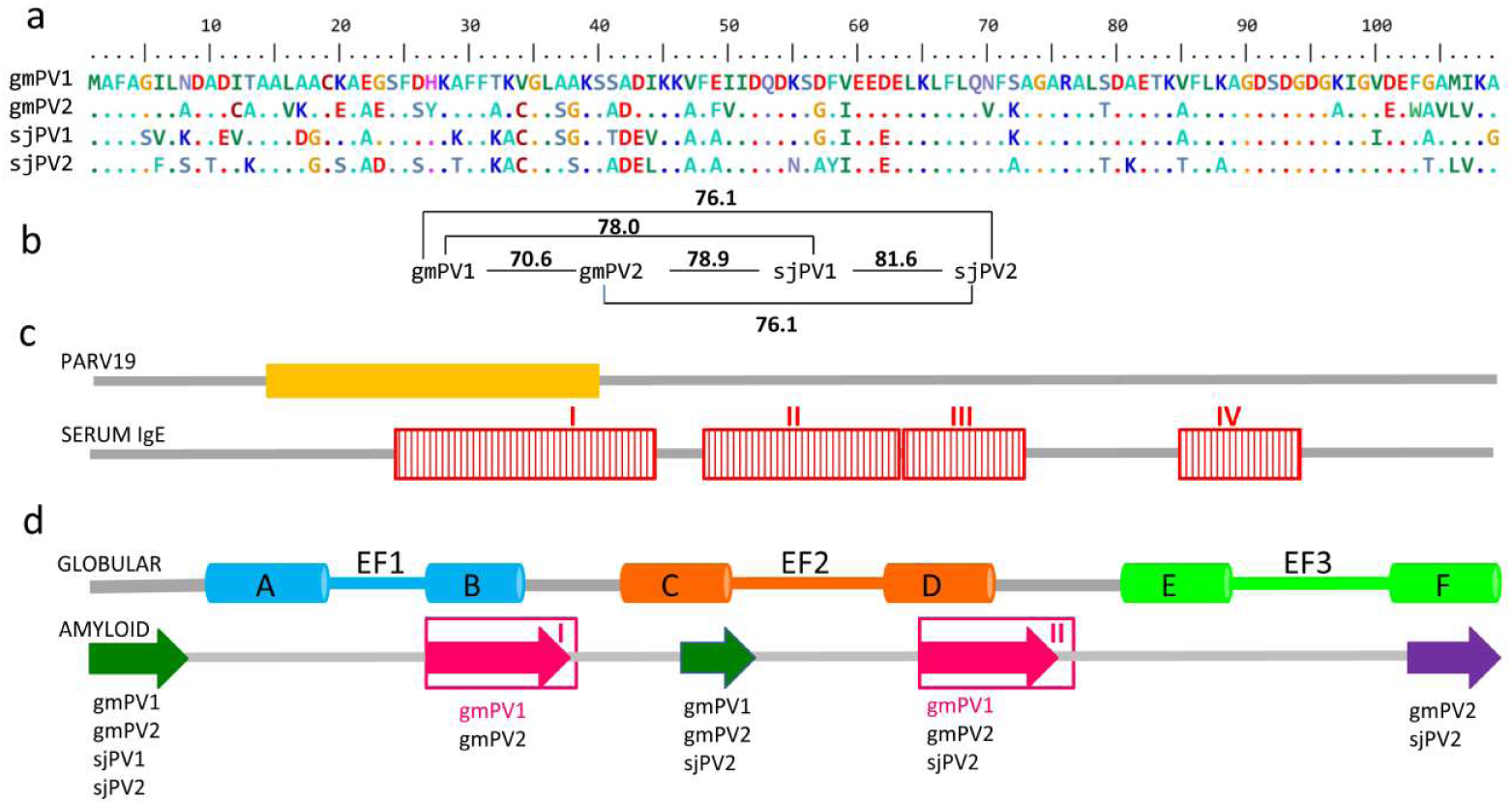
Sequences of the β-PV isoforms of *Gadus morhua* and *Scomber japonicus*. **(a)** Sequence alignment generated using BioEdit tools and their amino acid color table**. (b)** Pairwise identity calculated using BioEdit tools**. (c)** Relevant regions for PARV19 and serum IgE binding. The PARV19 epitope (orange rectangle) was predicted to include residues 13-39^**36**^. The displayed IgE binding epitopes (red rectangles) were determined using gmPV1-templated dodecapeptides^**25**^. **(d)** Segments involved in the globular and amyloid folds of β-PV. The globular fold contains three helix-loop-helix (AB, CD and EF) EF-hands, of which only CD and EF bind cations^**3,4,26, 43**^. Regions assembling into amyloids were predicted by AmylPred2, and they are depicted by arrows. Regions known to form amyloids in gmPV1 are depicted with pink arrows^**25,30**^.

### β-PV isoform abundance in fish muscles

To determine the relative abundance of the isoforms, we analyzed fish muscle extracts. For geographical reasons, *Scomber scombrus* expressing Sco s 1 (UniProtKB D3GME4), which shares 99% sequence homology with sjPV1, was taken as a model of mackerel (data not shown). Analysis of muscle extracts prepared in TBS by SDS-PAGE using gmPV1 as a standard for quantitation of the monomer band showed that the PV content (mg PV /g tissue) amounts to 3.1 ± 0.4 and 0.6 ± 0.2 in *Gadus morhua* and *Scomber scombrus*, respectively (**Fig. 2a**). Incubation of the extracts at 37 °C (intrinsic proteolysis allowance) for 15 min showed a 10 and 30% reduction in the PV monomer in *Gadus morhua* and *Scomber scombrus*, respectively, suggesting differences in the protease sensitivity of the PV and in the protease load of the extracts (**Fig. 2b**). Ultracentrifugation assays showed differences in the PV solubility between fish species extracts (**Fig. 2b**). Approximately 30% of the total *Gadus morhua* PV monomer partitioned in the pellet fraction, whereas the PV remained quantitatively soluble in *Scomber scombrus* extracts (**Fig. 2b**). Mass spectrometry showed that gmPV1 and sjPV1 were the most abundant forms in each fish species, representing approximately 85% of the total PV content, whereas gmPV2 and sjPV2 represented minor forms (**Fig. 2c**). Yet unknown PV isoforms such as one with a molecular weight of 11,784 Da were detected in *Scomber scombrus*.

**Figure 2.**
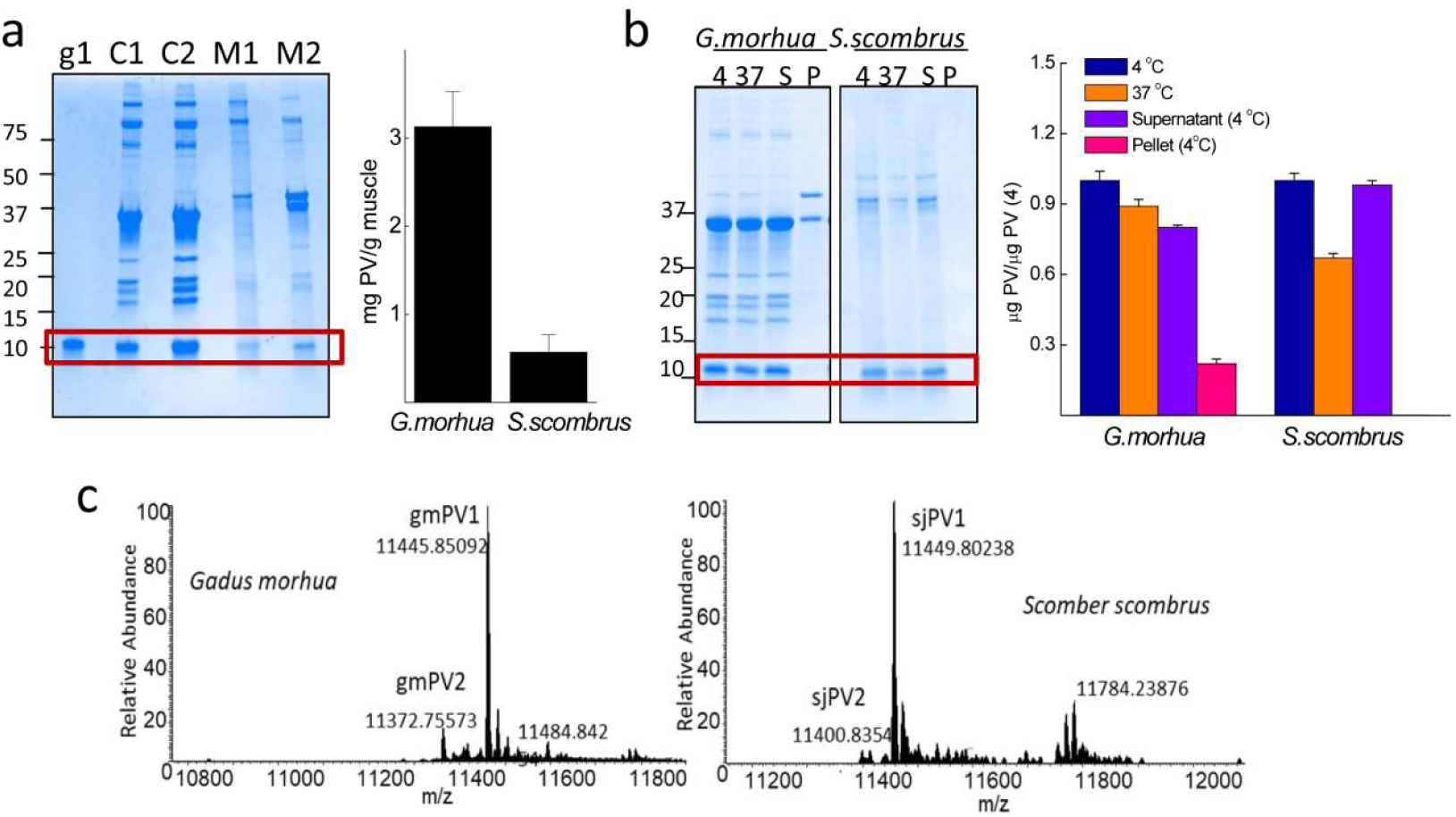
Total PV content and relative abundance of β-PV isoforms in *Gadus morhua* and *Scomber scombrus* muscles. **(a)** Typical Coomasie Blue-stained SDS-PAGE gel of (C1, C2) *Gadus morhua* and (M1,M2) *Scomber scombrus* muscle extracts and the PV content estimated from monomer band quantification. The protein load per lane was 5 μg for the extracts and 0.5 μg for gmPV1, which was used as a control. Numbers on the right side indicate the molecular weights of markers in kDa. (**b**) SDS-PAGE analysis of the intrinsic proteolysis and solubility of PV in muscle extracts. Freshly prepared extracts were (4) stored at 4°C, (37) heated for 15 min at 37°C, cooled at 4°C, and separated into soluble (S4) and insoluble (P4) fractions by ultracentrifugation. Numbers on the right side indicate the molecular weights of markers in kDa. **(c)** *Mr* determination for each of the different β-PV isoforms isolated from muscle extracts by FTICR-MS, considering the processing of M1 and the acetylation of A2^**39,52**^. The original gels of panels a and b are displayed in **supplementary Fig. S1.**

### Sequence-dependent features of the IgE interaction with β-PVs

To gain insight into the sequence factors involved in the interaction with IgE, the β-PV chains were denatured under reducing conditions and analyzed by immunoblot (**Fig. 3**). To allow signal analysis via antibody recognition, protein loading was first verified by Coomassie Blue staining using concentrated stocks (**Fig. 3a**). The reactivity of the denatured chains was first probed using the PARV19 monoclonal antibody, which is predicted to recognize the region of residues 13-39 and is often used for fish PV quantifications^**4,36,38,41**^. PARV19 recognizes the 11 kDa bands of β-PV monomers. For samples with similar protein loading, sjPV1 was the only isoform that exhibited PARV19 positivity (**Fig. 3b**). When the relative protein loading of sjPV1 was decreased by 10-fold, PARV19 also recognized gmPV1 and sjPV2 but failed to interact with gmPV2 (**Fig. 3b**). Screening of the gmPV2 sequence for unique substitutions in the region of residues 13-39 suggested C12-A13-V16-K17-E20-Y27-A33 as the group of residues impairing PARV19 recognition (**Fig. 1a**). It must be noted that differences in PARV19 recognition of β-PV isoforms have also been described for the *Evynnis japonica* chains^**38**^. Therefore, these and previous results preclude the use of PARV19 reactivity for β-PV complex quantifications. In fact, if used in muscle extracts, the obtained quantifications would have underestimated the contents for *Gadus morhua* (lack of recognition of gmPV2) and overestimated the amounts for *Scomber jap*o*nicus* (increased affinity of sjPV1).

**Figure 3.**
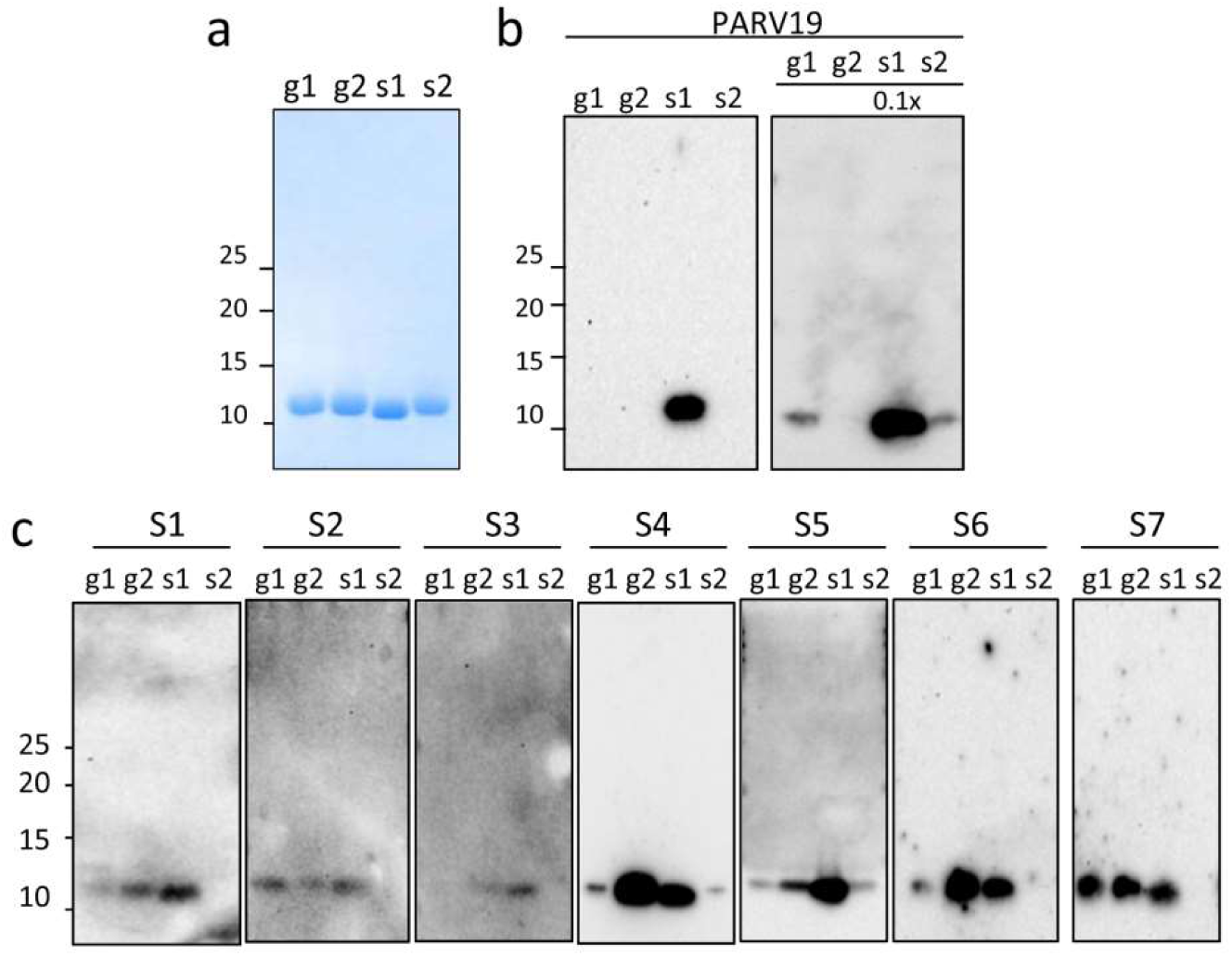
Immunoreactivity of the denatured state of β-PV: linear IgE epitopes. **(a)** Coomassie blue-stained SDS-PAGE gel of β-PV isoforms. **(b)** Western blots of β-PV isoforms probed with PARV19. The relative loading amount is depicted at the top. **(c)** Western blots of β-PV isoforms probed with different fish-allergic patient sera for IgE binding. Displayed blots correspond to membranes developed independently. Numbers on the right side of gels and membranes in panels a and b correspond to the molecular weights of markers in kDa. Full-length gels and blots are displayed in **supplementary Figs. 2S and 3S**.

Binding of IgE was then assayed using sera from patients with confirmed fish allergy (**Fig. 3c**). As shown by western blotting, serum IgE mainly recognized three isoforms, the interaction being significantly stronger with gmPV2 and sjPV1 than with gmPV1. Despite singularities, the pattern of recognition for gmPV1 was highly reproducible among distinct sera, supporting a general trend. The search of the residues uniquely shared by gmPV2 and sjPV1 yielded S37, G38, G57, K72, A85 and A104. Of them, residues 37 and 38 are contained in the linear major epitope (22-41 segment) described in both sjPV1 and gmPV1 using synthetic peptides and overlap the region that is considered to be the universal IgE epitope (**Fig. 1c**)^**3,4,22,25,27,43**^. Therefore, residues 37-38 seem to play a key role in dictating the affinity (SG>AA) and allowance (SG, AA vs AS) of the IgE interaction. Notwithstanding, unknown contributions from the unfolded states of the proteins, such as local refolding and aggregation upon SDS removal, cannot be ruled out.

### Conformational properties of β-PV isoforms

The effect of the sequence differences on the folding properties of the distinct β-PV isoforms was first analyzed using purified Ca^2+^-bound globular forms**. Fig. 4a** and **4b** show that freshly prepared Ca^2+^-bound proteins cooperatively folded with predominant α-helical secondary structures and hydrodynamic radii close to the R_H_^T^ of spherical monomers (1.6 ± 0.1 nm). These observations agree with the available 3D structures of gmPV1 and sjPV1^**26,43**^. However, despite these similarities, Ca^2+^-bound sjPV1 and gmPV2 unfolded at a significantly lower temperature than the other chains (**Fig. 4c**). It must be noted that sjPV1 and gmPV2 unfolding was largely irreversible based on the lack of recovery of the initial CD spectrum after the samples cooled from the highest temperature (**Fig. 4b**). Incubation of β-PV isoforms with proteinase K showed the complete degradation of sjPV1, the partial sensitivity of gmPV2 and the resistance of gmPV1 and sjPV2 (**Fig. 4d**). Since analysis of the sequences using the Expasy Peptide cutter tool showed that all isoforms contain more than 55 proteinase K cleavage sites along the chains, the sensitivity to protease digestion appears to inversely correlate with the globular fold thermal stability. When digestions were performed with pepsin at acidic pH, sjPV2 was fully digested, whereas the other proteins generated a resistant core (**Fig. 4d**).

**Figure 4.**
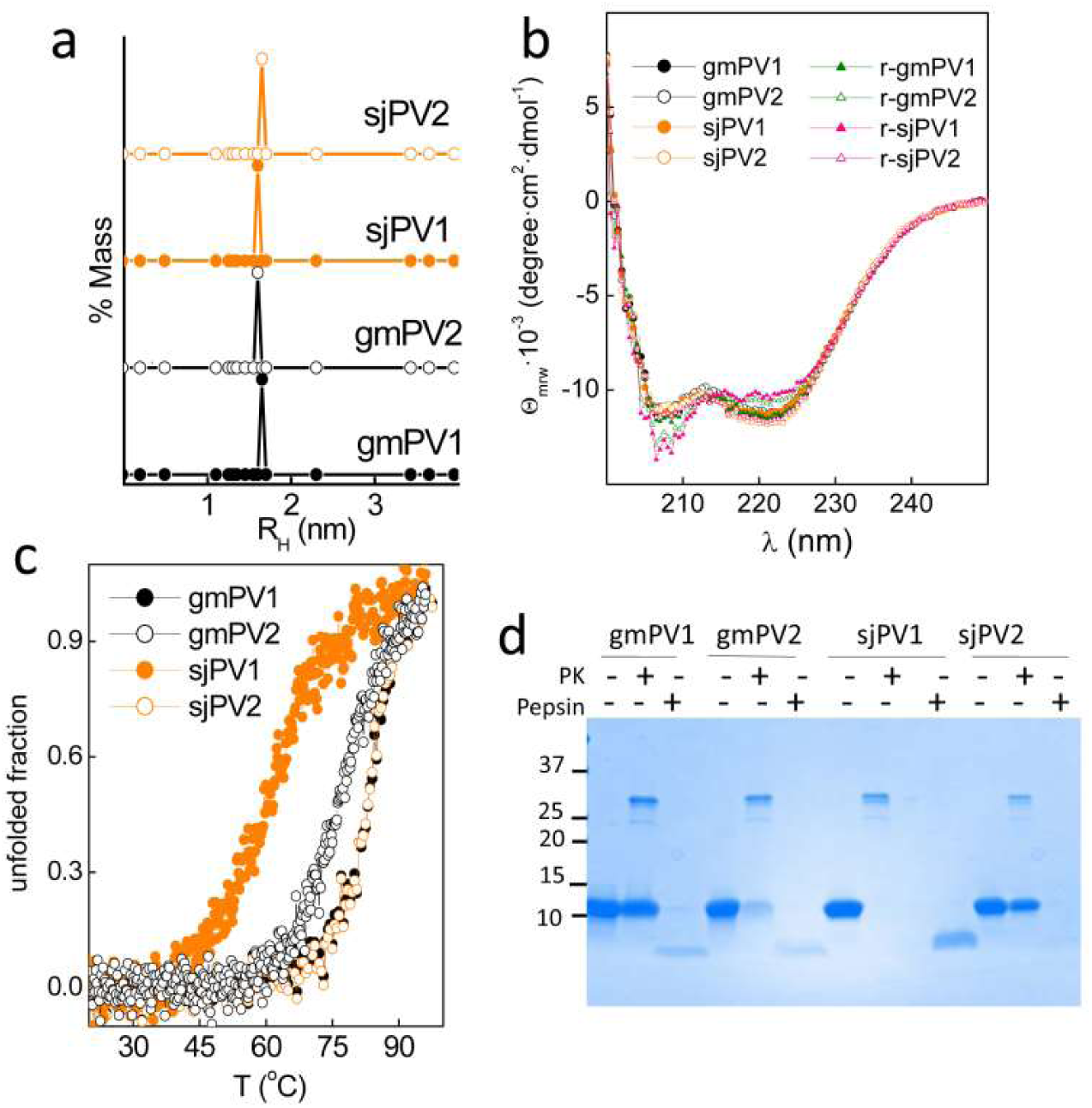
Conformational properties of the Ca^2+^-bound globular folds of the distinct β-PV isoforms. **(a)** DLS analysis of the association state and hydrodynamic features**. (b)** Secondary structure probed by far-UV CD spectra. Spectra were recorded before (circles) and after heating (triangles) at 95 °C. **(c)** Thermal denaturation monitored changes in θ_222_ as a function of temperature. Symbol key: •, gmPV1; •, gmPV2;, sjPV1; ○, sjPV2. (**d**) SDS-PAGE analysis of proteinase K and pepsin digestion products of β-PV isoforms. Digestions were performed using a 1/50 protease/β-PV ratio. The full-length gel is displayed in **supplementary Fig. 4S**.

Previous work showed that the removal of Ca^2+^ by either acidic pH or addition of chelates allowed the refolding of gmPV1 into fibrillary and insoluble β-sheet rich aggregates known as amyloids^**25,29**^. Since amyloid formation is a sequence- and concentration-dependent process, isoforms may display distinct competency. To evaluate the occurrence of amyloid assembly, β-PVs were incubated at 70 µM with 10 µM thioflavin T (ThT) under neutral (25 mM Tris-HCl pH 7.5, 50 mM NaCl, and 4 mM EDTA) and acidic (0.1 M Gly, pH 1.6) aggregation conditions, and the changes in the amyloid-probe fluorescence were monitored (**Fig. 5a**). The samples gmPV1, gmPV2 and sjPV2 exhibited a time-dependent increase in the fluorescence emission of ThT. In contrast, no changes were observed for sjPV1. Increasing the monomer concentration using increased incubation times (>150 h) and the inclusion of either 0.1 M GdnCl or 5 mM tris(2-carboxyethyl)phosphine (TCEP) in the reaction did not modify this aggregation trend (data not shown). Processes such as differential degradation were discarded from the analysis of the monomer mass conservation during the incubation period (**Fig. 5b**). Thus, these results indicated that the predicted amyloid-forming region in sjPV1 lacks functionality either intrinsically or due to an inhibitory effect of the C-terminal flanking region (**Fig. 1d**)^**46,47**^.

**Figure 5.**
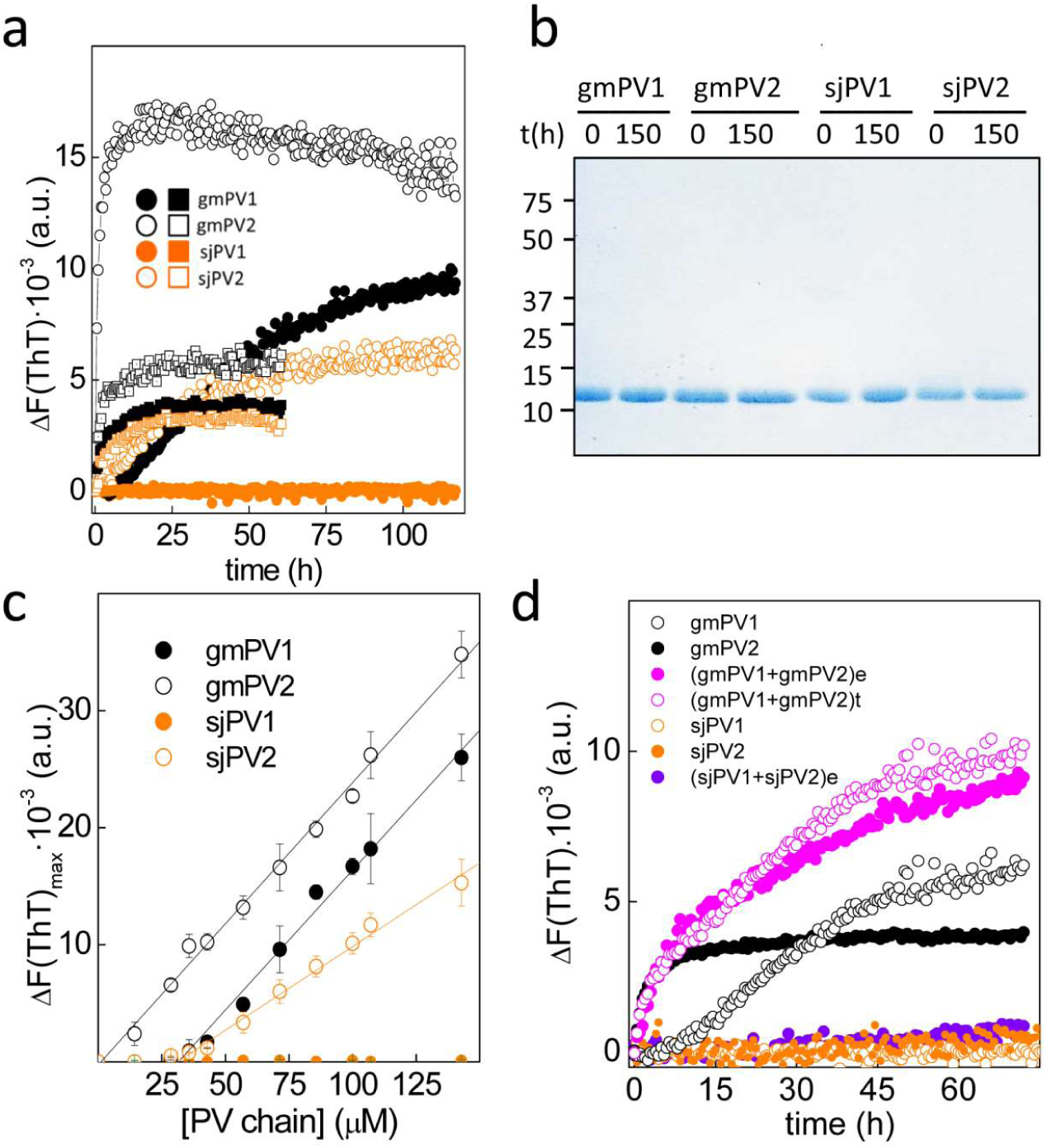
Amyloid formation by β-PV isoforms. **(a)** ThT-binding kinetics of gmPV1 (solid black symbols), gmPV2 (open black symbols), sjPV1 (orange solid symbols) and sjPV2 (orange open symbols) under acidic (square) and neutral (circle) aggregation conditions. **(b)** SDS-PAGE analysis of β-PV monomer conservation during amyloid formation at neutral pH. The full-length gel is displayed in **supplementary Fig. 5S. (c)** Intensity of ThT fluorescence as a function of the concentration of β-PV monomers. Values correspond to kinetics performed at pH 7.5 in the presence of EDTA. **(d)** Aggregation kinetics of 75 μM PV1 in the absence and presence of 15 μM PV2 (e) obtained experimentally in mixed reactions or (t) generated theoretically summing the traces of each of the chains.

Plots of the change in fluorescence intensity as a function of the monomer concentration show 31.1 μM, 1.1 μM and 31.1 μM as the critical concentrations for the aggregation of gmPV1, gmPV2 and sjPV2, respectively (**Fig. 5c**). These data indicated that gmPV2 aggregates at concentrations above 1.1 μM, whereas gmPV1 and sjPV2 require a 10-fold higher concentration for their aggregation. To better simulate the conditions in muscle extracts, the effect of PV2 (minor) on PV1 (major) aggregation was then studied. **Fig. 5d** shows that the experimental kinetic trace of an aggregation reaction containing 75 μM gmPV1 and 15 μM gmPV2 overlapped with a theoretical trace generated from the sum of the individual aggregation reactions. Additionally, the presence of 15 μM sjPV2 did not modify the non-aggregating pattern of sjPV1. These data supported the absence of coaggregation events.

CD spectroscopy confirmed the presence of β-sheet signatures in gmPV1, gmPV2 and sjPV2 insoluble aggregates (**Fig. 6a**). Despite this similarity, the use of anti-amyloid conformational antibodies revealed structural differences among the amyloid aggregates^**30**^. Thus, gmPV1 and gmPV2 aggregates contained amyloid-fibril epitopes (OC reactivity), whereas those formed by sjPV2 contained solely β-oligomer epitopes (A11 specificity) (**Fig. 6b**). These differences impact the protease resistance profile of the resulting aggregates. **Figure 6c** shows that exposure of the gmPV1 and gmPV2 OC-reactive aggregates to harsh proteolysis yielded pepsin- and proteinase K-resistant cores. In contrast, the A11-reactive sjPV2 aggregates were fully digested under similar conditions.

**Figure 6.**
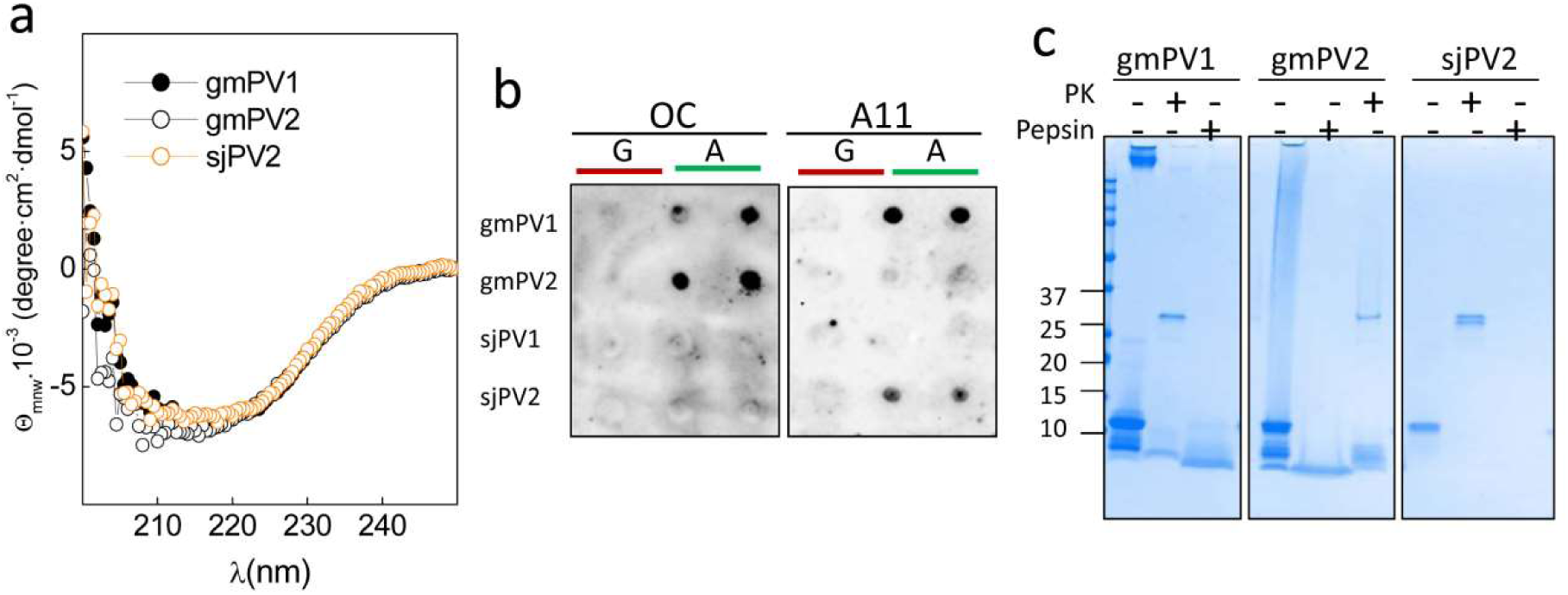
Features of the amyloid aggregates formed by β-PV isoforms. **(a)** Far-UV CD spectra of the insoluble aggregates of gmPV1, gmPV2 and sjPV2. **(b)** Dot blot analysis of (A) β-PV aggregates using amyloid-specific anti-fibril (OC) and anti-oligomer (A11) antibodies. (G) Ca^2+^-bound chains were included as controls. **(c)** Coomassie blue-stained SDS-PAGE gel of the harsh proteinase K and pepsin digestions of gmPV1, gmPV2 and sjPV2 amyloid aggregates. The original membranes and gels of panels b and c are displayed in **supplementary Fig. 6S**.

Together, these data showed that isoforms differ in the stability and protease sensitivity of the globular fold (sjPV1 and gmPV2 displayed thermal lability and protease sensitivity), in the capacity to form amyloids (sjPV1 is amyloid prone), in the concentration requirements for assembly (gmPV2<< gmPV1 and sjPV2) and in the conformation (OC vs A11 reactivity) and protease sensitivity of the aggregated state. Additionally, the coaggregation of isoforms from the same fish species is discarded as a possibility.

### Conformation-dependent traits of the interaction of IgE with β-PVs

To analyze and compare the IgE binding to the isoforms in the globular and amyloid folded states, we used a dot blot assay (**Fig. 7).** This assay avoids the interference of events such as precipitation, the dissociation of oligomers, pH-induced conformational changes and differential adherence during the ELISA coating step and allows the direct quantitation of protein loads by the Ponceau red staining of duplicates. As observed with the denatured states, serum IgE interacted the most with the Ca^2+^-bound globular monomers of gmPV2 and sjPV1. Preheating of Ca^2+^-bound globular monomers caused a signal reduction but preserved the trend of interaction of the unheated monomers. Removal of the bound Ca^2+^ from the monomers by the addition of EDTA in dilute solutions, both in the absence and presence of heat treatment, reproduced the previous recognition pattern, indicating that the allergenicity of the globular fold is likely governed by independent sequential Ca^2+^-binding epitopes. Furthermore, the absence of a significant effect of the preheating treatment on sjPV1 immunoreactivity and the lack of IgE recognition of sjPV2 suggested that the reported heat sensitivity of mackerel allergenicity might be linked to the lack of protease resistance of the major isoform^**36,42**^. On the other hand, the analysis of the IgE recognition of gmPV1, gmPV2 and sjPV2 amyloids showed immunoreactivity only in gmPV1 aggregates (**Fig. 7a,b**). Then, the amyloid fold appeared to be a regulator of IgE interaction, acting as an inhibitor for gmPV2 and as an amplifier for gmPV1, as previously reported^**25**^. The absence of a serum interaction for sjPV2 comparable with such interactions for the rest of the isoforms made this protein a potential hypoallergenic molecule.

**Figure 7.**
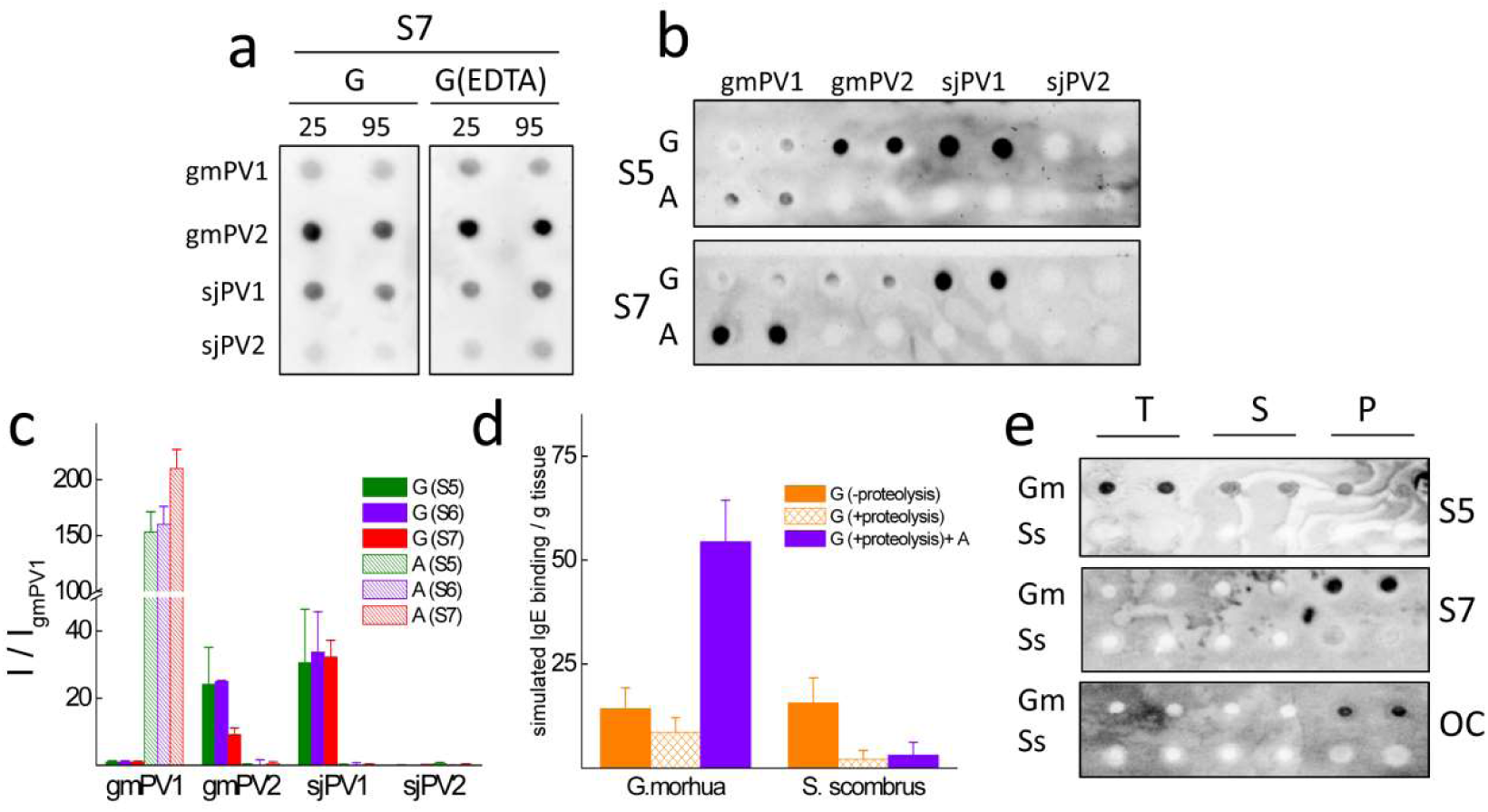
Comparative serum IgE interaction of β-PV isoform folds. **(a)** Typical dot blots of the effect of temperature treatment of β-PV on the recognition by the IgE present in sera from fish-allergic patients. Key: G, Ca^2+^-bound globular monomer; G(EDTA), Ca^2+^-bound globular monomer treated with 5 mM EDTA; 25 and 95 °C, temperatures at which samples were kept for 15 min. (**b**) Typical dot blots of the effect of the β-PV isoform fold on the recognition by the IgE present in sera from fish-allergic patients. Key: G, Ca^2+^-bound globular monomer; A, amyloid aggregate; Si, sera from distinct fish-allergic patients. (**c**) IgE binding intensity obtained for the different conformations of β-PV isoforms compared to that of the Ca^2+^-bound globular fold of gmPV1 for distinct sera. (**d**) Simulation of fish muscle IgE binding from the content and properties of component β-PV isoforms. Key: G (-proteolysis), globular folds under proteolysis inhibition; G (+proteolysis), globular folds with proteolysis; G (+proteolysis)+A, globular folds with proteolysis and amyloids. (**e**) Serum IgE and OC immunoreactivities of *Gadus morhu*a (Gm) and *Scomber scombrus* (Ss) muscle extracts (T) and their supernatant (S) and pellet (P) fractions. Blots were performed in duplicate using duplicated dots. All blots were loaded with 100 ng of protein per dot. The original membranes used in a, b and e panels are displayed in **supplementary Fig. 7S.**

### Reconstruction of the species-dependent IgE reactivity of fish muscle from PV isoform characteristics

To gain insights into the relative β-PV allergenic potency of fish muscle extracts, IgE binding was simulated as a linear combination of content and serum average immunoreactivity of the component isoforms for the different structural states (**Fig. 7c**). Under row experimental conditions and the impedance of aggregation and proteolysis events, the IgE binding signal would be provided by that of the distinct globular folds, with both fish species yielding similar levels. In the absence of protease inhibitors, sjPV1 and gmPV2 would be degraded, resulting in a relatively higher allergenicity for *Gadus morhua* than for *Scomber japonicus/scombrus*. In fact, the use of the degradation factors depicted in **Fig. 2b** (10 and 30% reduction in the PV monomer band in *Gadus morhua* and *Scomber scombrus*, respectively) showed a 1.3-fold higher PV allergenicity for *Gadus morhua* than for *Scomber japonicus/scombrus*. Furthermore, assuming that all the insoluble PV shown in **Fig. 2b** is amyloid folded, the PV IgE binding of *Gadus morhua* was 17-fold higher than that of *Scomber japonicus/scombrus*.

Although fish muscle extracts can contain allergens other than PV^**21,48**^, to experimentally test the simulation results, the immunoreactivity of the extracts and their soluble and insoluble fractions was analyzed by dot blot. **Fig. 7d** shows that serum IgE recognized *Gadus morhua* as having the best immunoreactivity, largely due to the anti-amyloid fibrils OC antibody-reactive insoluble fraction.

## DISCUSSION

Fish allergy constitutes a good model to study food allergens *a priori* since 70-95% of its cases are elicited by the fish muscle protein β-PV^**1,3,21,49**^. Despite this apparent simplicity, β-PV is not a single entity but rather a complex collection of isoforms in each fish species tailored to accomplish muscle relaxation rate modulation during swimming^**50**^. In addition to the physiological implications, the sequence differences among β-PV isoforms define their capacity to act in sensitized individuals as IgE ligands and consequently to trigger IgE-mediated pathogenic events^**1,2,3,4**^. For the muscles of fish species such as *Gadus morhua* and *Scomber japonicus*, which were taken as models of high and intermediate β-PV content, sequence banks provide two major sets of isoforms with pairwise sequence identities that are above 70%. Here, we have identified and quantified the isoforms in fish muscles, addressed how the diversity impacts the recognition by serum IgE from fish allergic patients using the three major conformations of β-PVs and reconstructed the allergenic potency combining the content and the properties.

Quantitative analysis showed notable differences in the relative abundance of these isoforms in muscle extracts, such as those used in allergy diagnosis tests. For 100 g muscle samples, *Gadus morhua* will contain approximately 263.5 and 46.5 mg of gmPV1 and gmPV2, respectively, whereas a *Scomber* filet of similar weight will contain approximately 51 and 9 mg of sjPV1 and sjPV2, respectively. If all isoforms would interact similarly with serum IgE, then the *Gadus morhua* having higher allergenicity than *Scomber japonicus* could have been solely explained in terms of allergen dose, as previously proposed^**4,13,14,51**^. However, sequence and concentration differences impact folding, stability and interaction properties. Of the four β-PV isoforms studied, only gmPV2 and sjPV1, with a similar abundance, displayed optimal linear epitopes, resulting in the best IgE recognition of both the unfolded and globular states. However, their high allergenicity was counteracted by the low thermal stability and protease sensitivity of their globular folds and the lack of amyloid formation or the generation of non-immunoreactive amyloids. Then, their contribution to IgE binding depends largely on the intrinsic proteolytic extent of extracts. On the other hand, gmPV1 and sjPV2 both formed temperature-stable and protease-resistant globular folds and exhibited amyloid assembly capacity. Of these chains, sjPV2 was not recognized by serum IgE in any of the structural states considered, and gmPV1, displaying a weak interaction when denatured and globularly folded, exhibited the same level of IgE recognition as gmPV2 and sjPV1 upon amyloid assembly, which is ensured by its increased abundance. Indeed, anti-amyloid fibril OC-reactive species were detected in the insoluble fractions of *Gadus morhua* extracts. In sum, *Gadus morhua* muscles are equipped with an isoform combination and content that ensures IgE recognition of all PV folds. In contrast, the *Scomber japonicus* allergenic load is under proteolytic control. It must be noted that immunological benefits, if any, arising from the hypoallergenic sjPV2 will be reduced based on the protease sensitivity of its aggregates.

Previous tolerance studies using open food challenges have reported lower eliciting doses and higher symptom severity for cod than for mackerel^**8,10,11**^. These findings were interpreted in terms of cod being the primary sensitizer due to the dietary traditions of the group under study, and contributions from unknown differences in β-PV from cod and mackerel were also considered. Here, we have demonstrated using sera from pediatric patients sharing hake and megrim as offending fish species that these differences exist and that the obtained IgE reactivity pattern supports a lower probability of tolerance for *Gadus morhua* than for *Scomber japonicus* in fish-sensitized patients. Notwithstanding, it must be noted that increasing the number of fish species and the use of a patient cohort with a varying geographical distribution is needed to strengthen the overall conclusion.

Contribution of both content and isoform composition to the relevance of β-PV cross-reactivity underlines the importance of the definition of common methodologies and actions. A reevaluation of the β-PV isoform composition and content of fish species using actual proteomic tools may be helpful for resolving this issue. The development of novel monoclonal antibodies universally recognizing β-PV will prevent the requirement of correction factors for the use of PARV19 in β-PV quantifications^**51**^. Additionally, the protease sensitivity and aggregation capacity of the distinct isoforms must be taken into account as a source of IgE reactivity variability. In this sense, a recent analysis has shown a large variation in the β-PV content in the commercial fish extracts used for skin prick testing, suggesting the possible interference of uncontrolled intrinsic proteolysis ^**48**^. In other grounds, the clinical determination of fish-specific IgE levels is performed using a variety of extracts lacking accurate definitions of β-PV isoforms, content and properties. Our results suggest that expanding the extract conditions to include all possible isoform conformations and their properties may provide a better understanding of the clinically relevant IgE reactivity.

## MATERIALS AND METHODS

### Ethics statement

All experimental protocols and methods were performed in accordance with the Hospital Universitario La Paz and the Consejo Superior de Investigaciones Científicas guidelines. Approval from the Ethics Committee of Hospital Universitario La Paz (PI1950) and the written informed consents of the patient parents were obtained.

### Patient sera

Serum samples were obtained from fish-allergic patients from the Allergy Service of the Hospital Universitario La Paz (Madrid, Spain) following the Ethics Committee of the Hospital Universitario La Paz guidelines. Serum samples from 7 fish-allergic patients (mean age: 8.4 years, 5 boys) with specific IgE antibodies to cod and tuna PV and volume availability (at least 4 ml) were selected. All patients had a history of symptoms suggestive of immediate hypersensitivity elicited by eating fish, specific IgE antibodies to fish as determined by CAP-System FEIA™ (Thermo Fisher, USA), and positive skin prick tests to distinct fish extracts: cod and tuna (1000 IC/mL, Alyostal, Stallergenes, France); swordfish and salmon (1 mg/mL, Bial Aristegui, Spain); and hake (1.25 mg protein/mL) and megrim (1 mg/mL, Laboratorios LETI S.L. Spain) (**Supplementary Table S1 online**).

### Fish muscle extract preparation and processing for total β-PV quantification, β-PV isoform purification and analysis by Fourier transform ion cyclotron resonance mass spectrometry (FTICR-MS)

Fish belonging to *Gadus morhua* and *Scomber scombrus* were purchased from local markets and genetically identified using a fish ID Kit (Bionostra S.L., Spain). For total PV quantifications, 0.5 g of muscle was homogenized in 1 mL of 25 mM Tris-HCl pH 7.4, 137 mM NaCl, 2.7 mM KCl (TBS) using an OMNI TH International homogenizer (2 cycles of 10 strokes in ice), followed by centrifugation at 4,000 x g for 5 min at 4 °C. Supernatants were collected, and their protein content was determined by the Bradford assay (see below). The soluble and insoluble fractions of extracts were obtained by 1 h ultracentrifugation at 100,000 x g at 4 °C using an Optima Tm^Max^ ultracentrifuge (Beckman, USA). Samples containing 10 μg of total protein were analyzed by SDS-PAGE and Coomassie Blue staining (see below). The amount of PV was determined from the band of the 11-12 kDa monomer using gmPV1 as a standard for quantification. Samples were analyzed using two distinct extracts in duplicate. For isoform analysis, the sarcoplasmic protein extraction was carried out by homogenizing 1 g of white muscle in 1 mL of MilliQ water for 30 s on ice in a sonicator device (IKA-Werke, Germany) as described^**39**^. β-PV extracts were acidified with 0.1% formic acid in 50% acetonitrile and analyzed using an LTQ FT Ultra mass spectrometer (Thermo Fisher Scientific, USA). For mass spectrometry, 50 µL of sample was analyzed by direct infusion at a flow rate of 1 µL/min. Ions were generated by positive ion mode using 4.5 kV for spray voltage. Proteins were analyzed with full-scan MS in the positive mode from 400 to 1,200 *m/z* applying 100,000 full widths at half mass of mass resolution. Data were collected for 5 min. The obtained signals were deconvoluted within XCalibur software (Thermo Fisher Scientific, USA) using the Xtract feature in the Qualbrowser program. All samples were analyzed in triplicate.

### Production of β-PV isoforms

Synthetic genes corresponding to the sequences of Gad m 1.0202 (gmPV1, A5I874) and Gad m 1.0102 (gmPV2, A5I873) from *Gadus morhua* (Atlantic cod) and Sco j 1 (sjPV1, Q3C2C3 and sjPV2, G9I591) from *Scomber japonicus* (chub mackerel) were obtained from Genscript (USA) in pET15b constructs. All proteins were produced in *Escherichia coli* BLD21 (DE3), isolated and purified as described previously ^**25,45**^. N-terminal His-tag removal was performed using a Thrombin CleanCleave™ kit as recommended by the manufacturer (Sigma-Aldrich, USA). Before their use, protein solutions were extensively dialyzed against 5 mM HEPES pH 7.5 containing 0.1 mM CaCl_2_, concentrated using 10 kDa-pore size Amicon Ultra-15 filters and centrifuged at 16,000 x *g* for 20 min at 4 °C to remove any possible aggregate.

### Formation of β-PV aggregated folds

The kinetics of amyloid formation was followed using a 96-well format Thioflavin T (ThT) binding assay. Cleared protein stocks (0.2 mM protein in 5 mM HEPES, pH 7.5, and 0.1 mM CaCl_2_) were diluted with distinct buffers to 14-150 µM protein and supplemented with 10 μM ThT. The buffers for the reactions were 25 mM Tris-HCl, pH 7.5, 0.1 M NaCl containing 4 mM either EDTA or CaCl_2_ (control for no aggregation) and 50 mM Gly, pH 1.6. Assays were initiated by placing the sealed 96-well plate at 37°C in a POLARstar (BMG Labtech, Germany) microplate reader. The ThT fluorescence was measured through the bottom of the plate every 30 min with a 450 nm excitation filter and a 480 nm emission filter. All measurements were performed in duplicate, and the experiments were repeated twice with different protein batches. When required, aggregates were harvested from the reaction mixtures by a 100,000 x g centrifugation for 1 h using an Optima Tm^Max^ ultracentrifuge (Beckman, USA). The resulting pellet and supernatant fractions were used for protein concentration determination. Before use, pellet suspensions were sonicated in a bath sonicator for 5 min.

### Dynamic light scattering

Dynamic light scattering (DLS) measurements were performed with a DynaPro spectroscatter (Wyatt Technology, USA) using 100 µM protein solutions as described previously^**25,29**^.

### Circular dichroism Spectroscopy

Circular dichroism (CD) measurements were performed with a Jasco J-820 spectropolarimeter (Jasco Inc., USA) as described previously^**25,29**^.

### Protease digestions

Solutions of globular monomers at 4 mg/ml in 5 mM HEPES, pH 7.5, and 0.1 mM CaCl_2_ were either left at 25 °C or heated at 95 °C for 15 min and then cooled to 25 °C for 10 min. Heated and unheated globular monomer samples were then diluted in either 50 mM Tris-HCl, pH 7.5, containing 0.1 M NaCl or 0.1 M Gly, pH 1.6. Similarly, aggregated protein samples at 4 mg/ml in 5 mM HEPES, pH 7.5, were diluted in either 50 mM Tris-HCl, pH 7.5, containing 0.1 M NaCl or 0.1 M Gly, pH 1.6. Solutions at pH 7.5 were incubated for 30 min in the absence and presence of proteinase K (Promega, 1:50 protease:isoform) at 37 °C, and the reactions were stopped with phenylmethylsulfonyl fluoride (Sigma-Aldrich, 5 mM final concentration). Solutions at pH 1.6 were incubated for 30 min in the absence and presence of pepsin (Promega, 1:50 protease:isoform) at 37 °C, and the reactions were stopped with 0.2 M Tris-HCl, pH 8.5. Reaction products were analyzed by SDS-PAGE.

### SDS-PAGE and Western blotting

Protein solutions were diluted in Laemmli sample buffer (2x) containing 2-mercaptoethanol, heated for 10 min at 100 °C and loaded on a 15% tris–glycine gel (10 µg protein/lane for Coomassie Brilliant Blue visualization; 0.2 µg protein/lane for immunoblotting). Precision Plus protein standards (Bio-Rad, USA) were used as standards for the molecular weights. Electrophoresis was carried out using a Mini-PROTEAN^®^ (Bio-Rad, USA) system at 100 V. For SDS-PAGE, proteins were visualized by Coomassie Brilliant Blue R-250 (Bio-Rad, USA) staining. For Western blotting, proteins were electrotransferred to PVDF membranes at 150 V for 1 h using prechilled 25 mM Tris, 192 mM glycine, pH 8.3, and 10% methanol transfer buffer in a Mini Trans-Blot^®^ Cell. Membranes were immersed in methanol, allowed to air dry and stored at −20 °C until use. For development, membranes were first blocked for 1 h in TBST-BSA (25 mM Tris-HCl pH 7.4, 137 mM NaCl, 2.7 mM KCl, 0.5% Tween-20, and 1% BSA). Immunodetection was performed by a 2 h incubation with PARV19 (Sigma-Aldrich, 1:1000 dilution) and sera from patients allergic to fish (1:10 dilution) prepared in TBST-BSA. After 3 x 10-minute washes, membranes were incubated for 1 h with horseradish peroxidase-labeled goat anti-mouse IgG (Sigma-Aldrich, 1:5000 dilution) or mouse monoclonal B3102E8 anti-human IgE (Abcam, 1:2000 dilution). After several washes with TBST, the signal was developed using the ECL-Clarity Western-blotting reagent (Bio-Rad, USA) and detected with a ChemiDoc XRS. When required, quantifications were performed using Image Lab software tools. For analysis, the signals in intensity/μg protein from at least two independent membranes and two sequential exposition times were referred to that of gmPV1 and averaged.

### Dot blot assays

Aliquots containing 15-150 ng of protein were spotted in duplicate on a nitrocellulose membrane and blocked for 1 h in TBST containing 1% BSA. Detection was performed by a 2-h incubation with the anti-amyloid fibril OC antibody (AB2286 Merck Millipore, 1:2000 dilution), anti-amyloid oligomer A11 antibody (AB9234 Merck Millipore, 1:2000 dilution) and sera from patients allergic to fish (1:10 dilution) prepared in TBST-1% BSA and a 30-min incubation with horseradish peroxidase-labeled mouse monoclonal B3102E8 anti-human IgE (Abcam, 1:2000 dilution) or goat anti-rabbit IgG (Sigma-Aldrich, 1:5000 dilution). The signal was developed by the ECL-Clarity Western blotting reagent (Bio-Rad) and detected with a ChemiDoc XRS. Quantifications and analysis were performed as before using the signals from gmPV1 as normalization factors.

### Protein quantification

Protein concentrations were determined using a Quick Start™ Bradford assay kit (Bio-Rad, USA), BSA standards and absorbance readings at 595 nm.

### Simulation of PV IgE binding in fish muscle extracts

Simulation of the IgE binding (IgEB) in muscle extracts was performed using a linear combination of the weighted contributions of the component isoforms in the different structural states as follows:

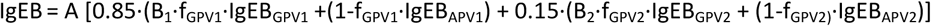

where A is the PV content in mg PV/ g tissue, f_GPVi_ is the fraction of the globular fold of the isoform i, B_i_ is the coefficient measuring the extent of intrinsic proteolytic degradation, and IgEB_GPVi_ and IgEB_APVi_ are the average intensities of the IgE binding of the globular and amyloid folds of the isoform i, respectively.

## ACKNOWLEDGEMENTS

This work was supported by grants from the Spanish AEI/EU-FEDER SAF2014-52661-C3 (MG), BFU2015-72271-EXP (MG), Angulas Aguinaga (MG, RRP) contract and the Ramón Areces Foundation (MC). MC is a Ramón y Cajal fellow. Laura Montoya and Rosa Sánchez are acknowledged for their technical support. The funders had no role in study design, data collection and analysis, decision to publish or preparation of the manuscript.

## AUTHOR CONTRIBUTIONS

MG and MC conceived and designed the experiments; RPT and MC conducted the experiments; MP, SQ and RRP obtained and characterized reagents; RPT, MC and MG analyzed the results; MG and MC drafted the manuscript. All authors reviewed the manuscript and contributed critical comments.

## COMPETING INTERESTS STATEMENT

Authors declare no competing interests.

