## Supplemental Information for "Reconstruction of fish allergenicity from the content and structural traits of the component β-parvalbumin isoforms"

[1]Insto Química-Física "Rocasolano", Consejo Superior de Investigaciones Científicas, 28006 Madrid, Spain.
[2]Insto Investigaciones Marinas, Consejo Superior de Investigaciones Científicas, 36208 Vigo, Spain
[3]Dpto de Alergología, Hospital Universitario La Paz, 28046 Madrid, Spain
[4]Insto de Investigación Hospital Universitario La Paz (IdiPaz), 28046 Madrid, Spain

### Supplementary Material

**Table 1**. Clinical data of the patients allergic to fish and their sera features.

| Patient | Age (years) | sex | Symptoms after fish ingestion[a] | Ofended fish[b] | Other allergies | SPT[c] (mm) Cod / Tuna | Total IgE (kU/l) | sIgE (kU/l) Cod / Tuna |
|---|---|---|---|---|---|---|---|---|
| S1 | 9.1 | M | U, OAS | Hake, tuna | Seafood[d], treenuts | 12 / 8 | 1,020 | 13.9 / 7.24 |
| S2 | 10.2 | F | U, AE, OAS, V | Hake, megrim | Fruits | 5 / 4.5 | 851 | 8.42 / 1.68 |
| S3 | 10.2 | M | V | Hake, cod, megrim | Egg, seafood, legumes, treenuts | 6 / 0 | 2,223 | 8.03 / 2.04 |
| S4 | 4.6 | M | AE, U | Hake | Egg | 15 / 12.5 | 901 | 15.3 / 1.83 |
| S5 | 7.8 | M | OAS, U | Hake, megrim | Egg, seafood, legumes, treenuts | 25.5 / 9.0 | 278 | 92.7 / 31.2 |
| S6 | 10 | F | AX | Hake | Seafood | 7.5 / 6.5 | 38.8 | 5.01 / 1.34 |
| S7 | 7.2 | M | OAS | Hake, megrim | Egg, nuts, legumes | 13 / 5.5 | 1,382 | 25.7 / 8.72 |

[a] Abbreviations correspond to: AE: angioedema; OAS: oral allergy syndrome; U: urticaria.

[b] Fish species causing initial symptomatology upon oral exposure.

[c] Mean diameter

[d] Molluscs and crustaceans.

**Figure 1S** : Original gels of Fig. 2a and 2b.

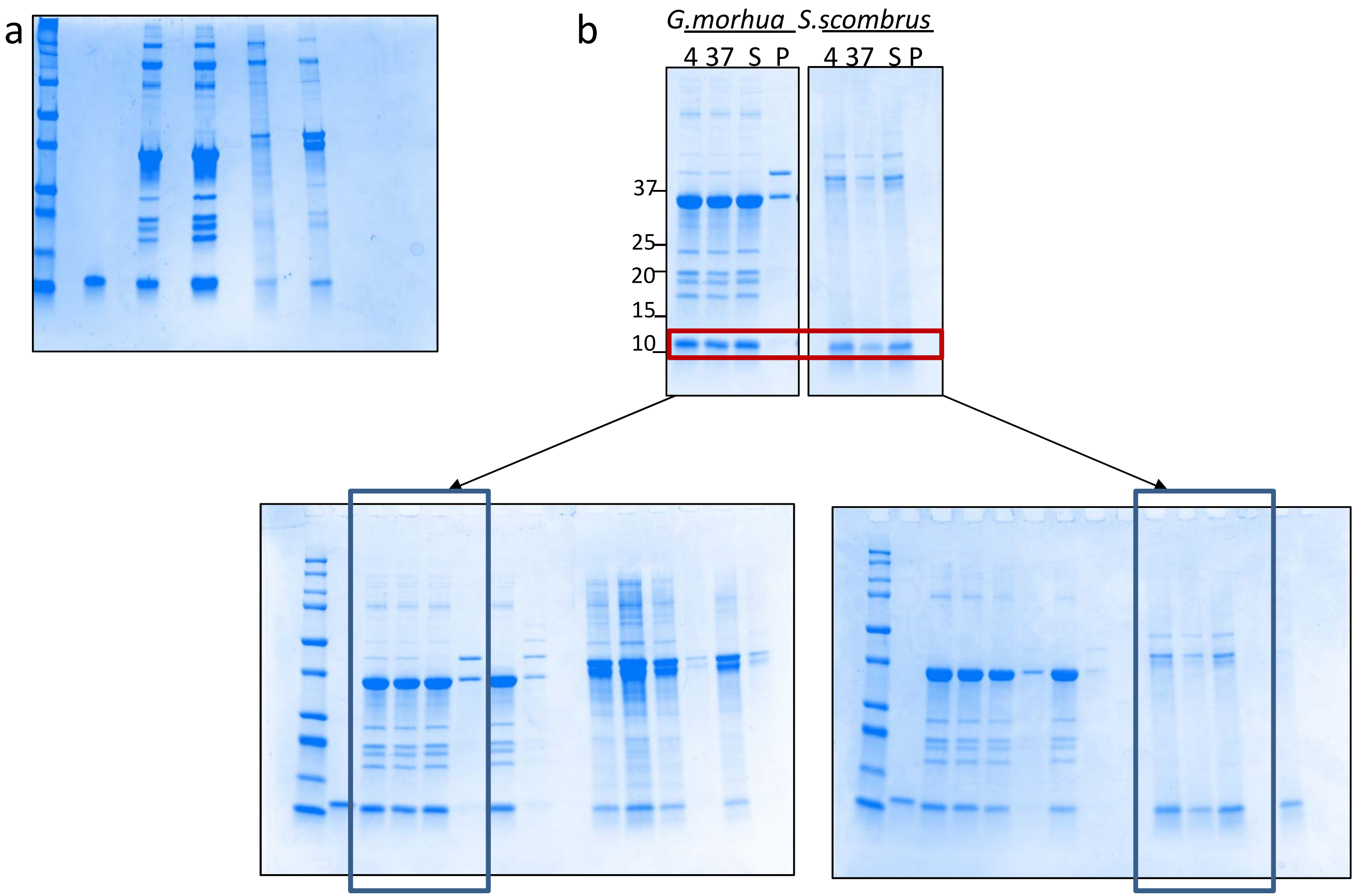

**Figure 2S** : Original gels and membranes of Fig. 3a and 3b.

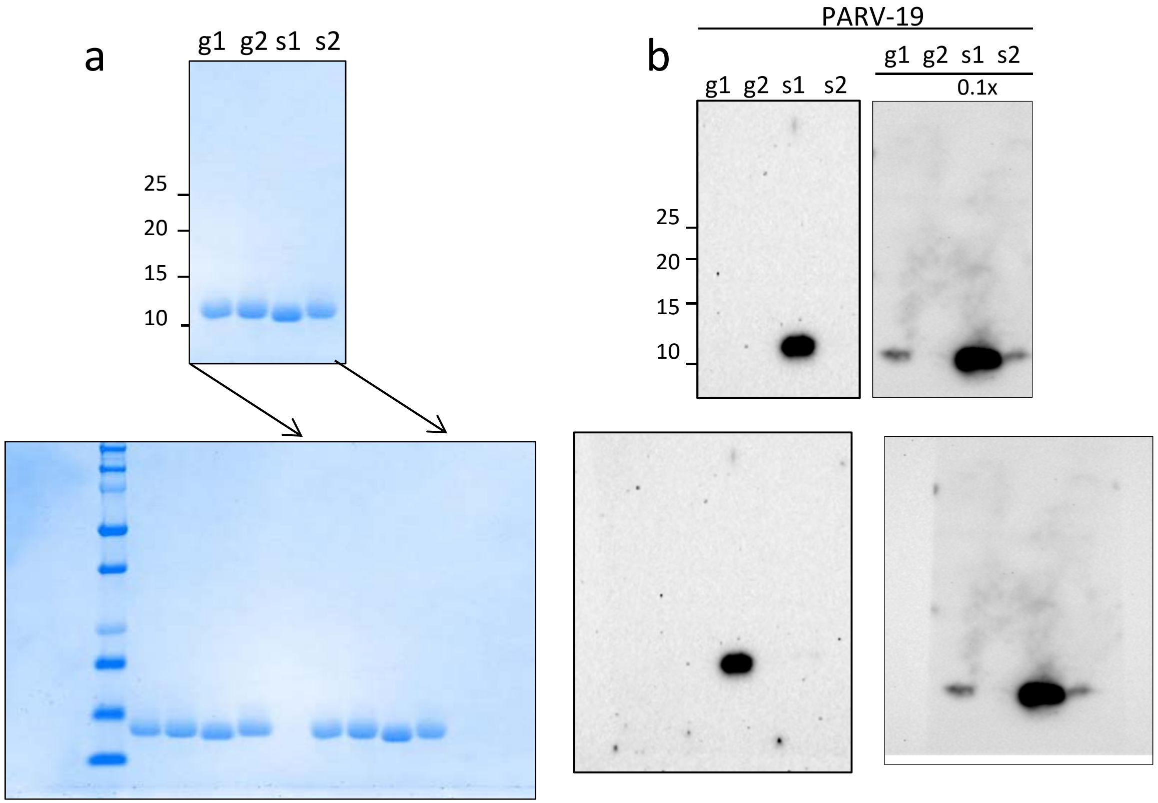

**Figure 3S** : Original blots used in Fig. 3c.

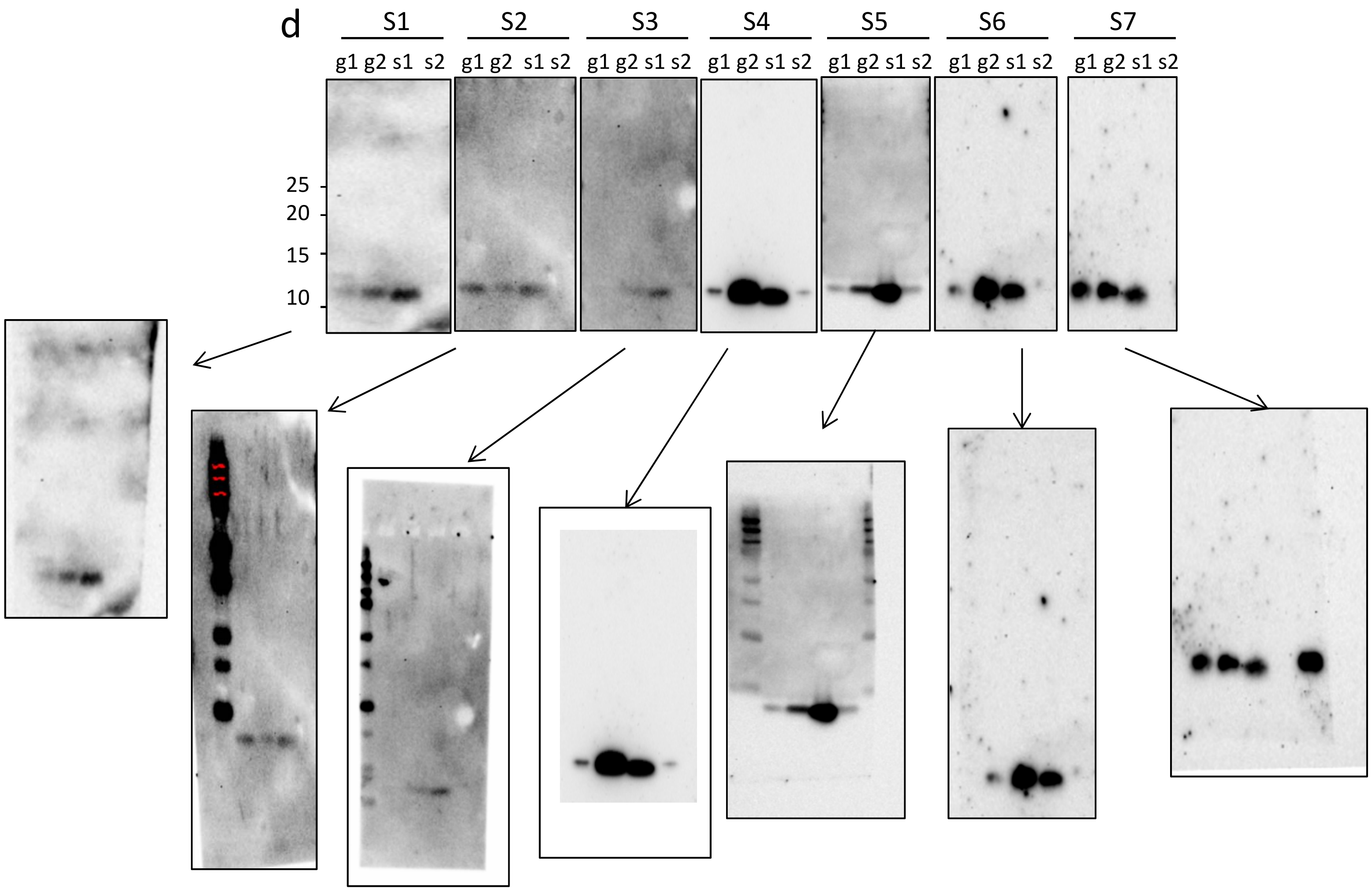

**Figure 4S** : Full length gel of Fig. 4d.

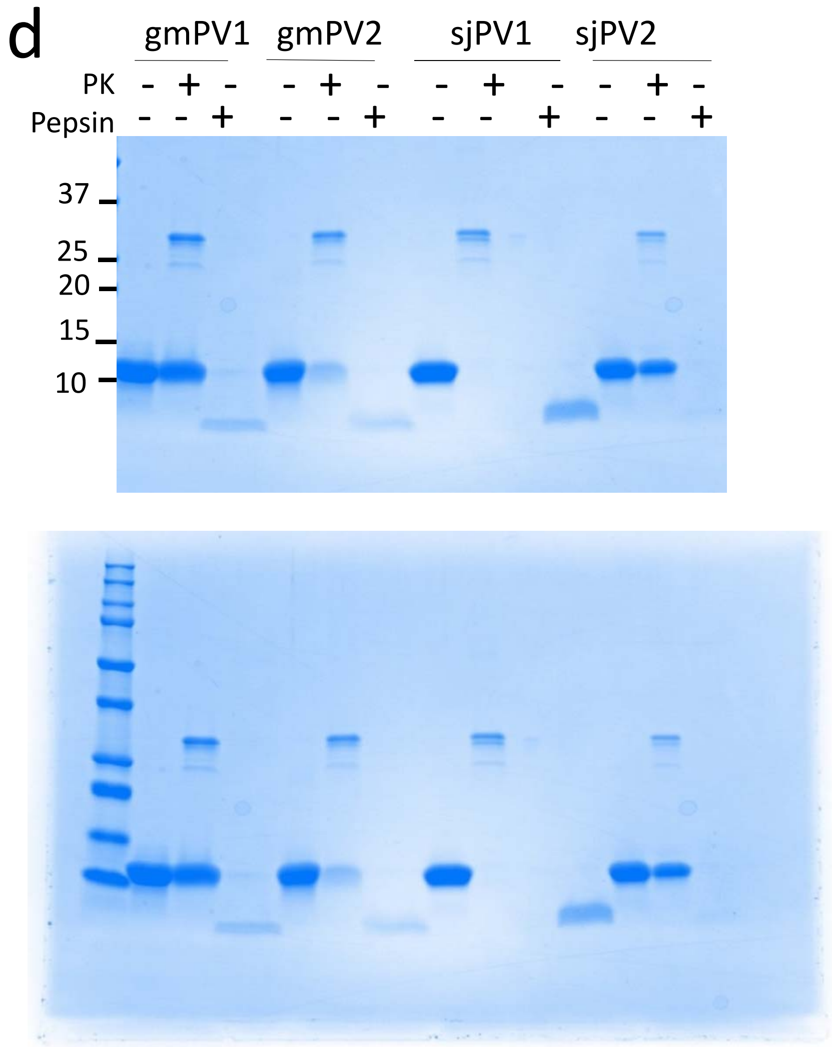

**Figure 5S** : Full-length gel of Fig. 5b.

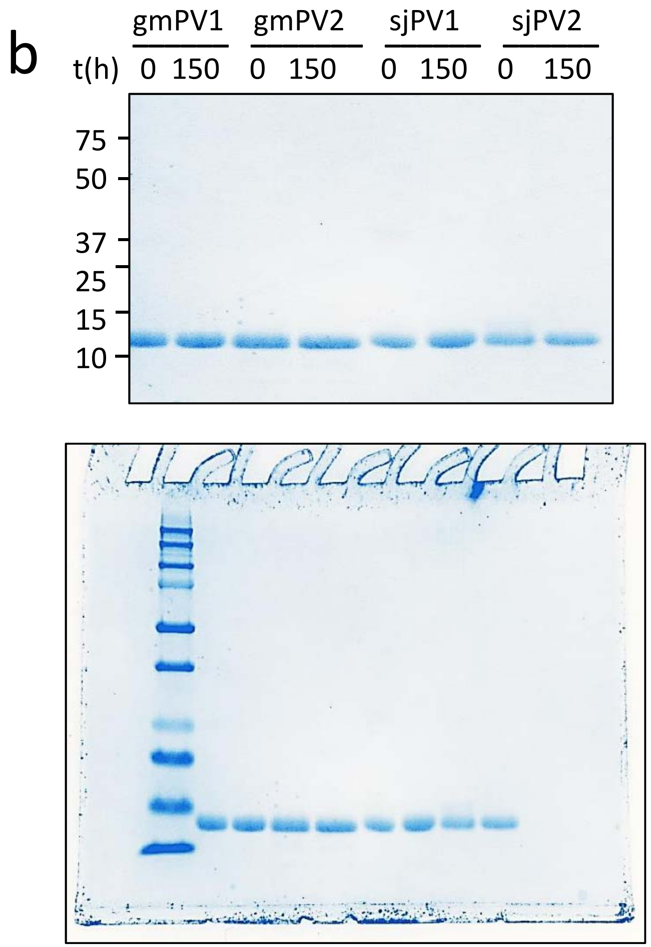

**Figure 6S** : Original membranes of Fig. 6b and 6c

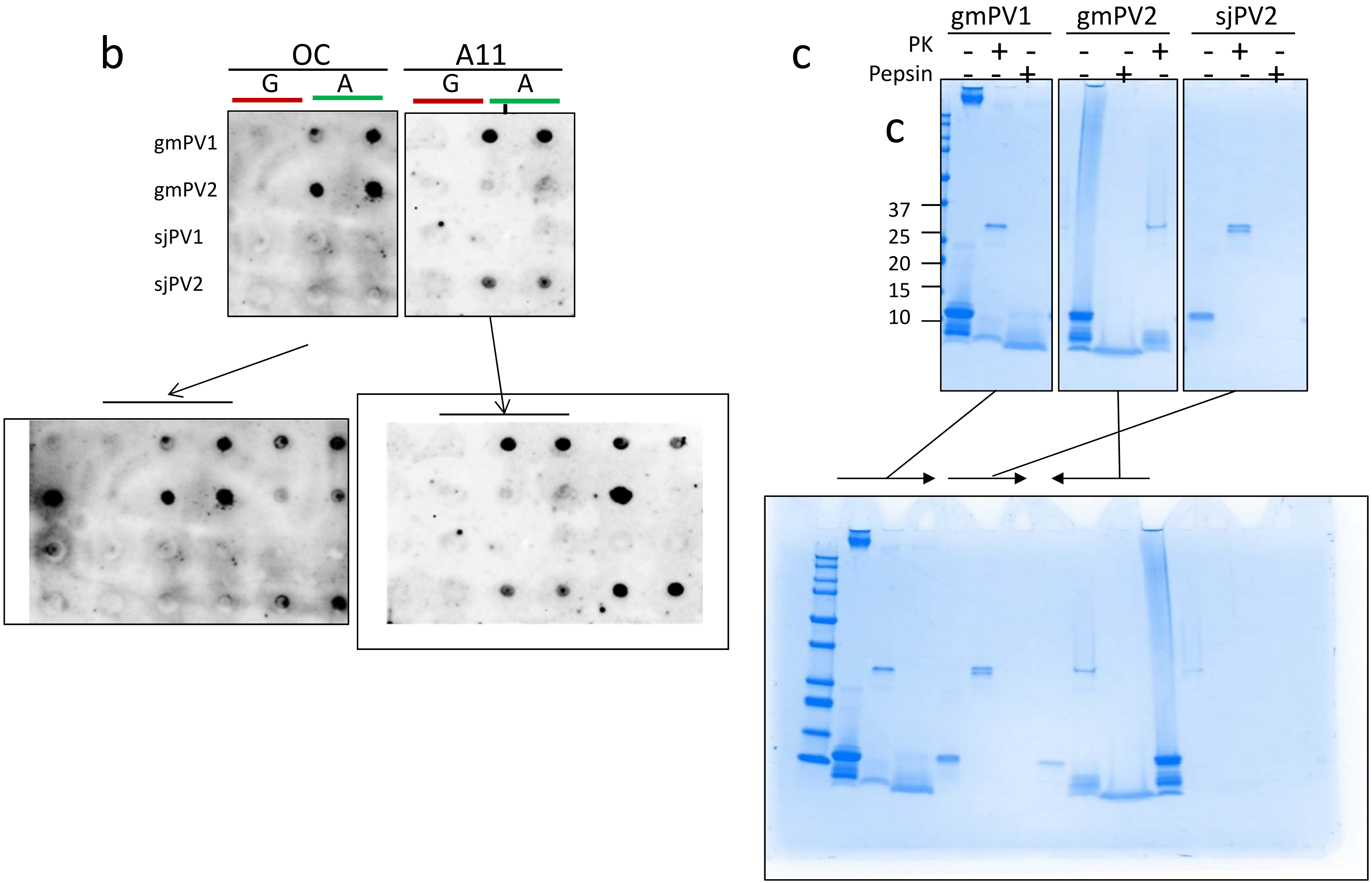

**Figure 7S** : Full size membranes of Fig. 7a, 7b and 7e.

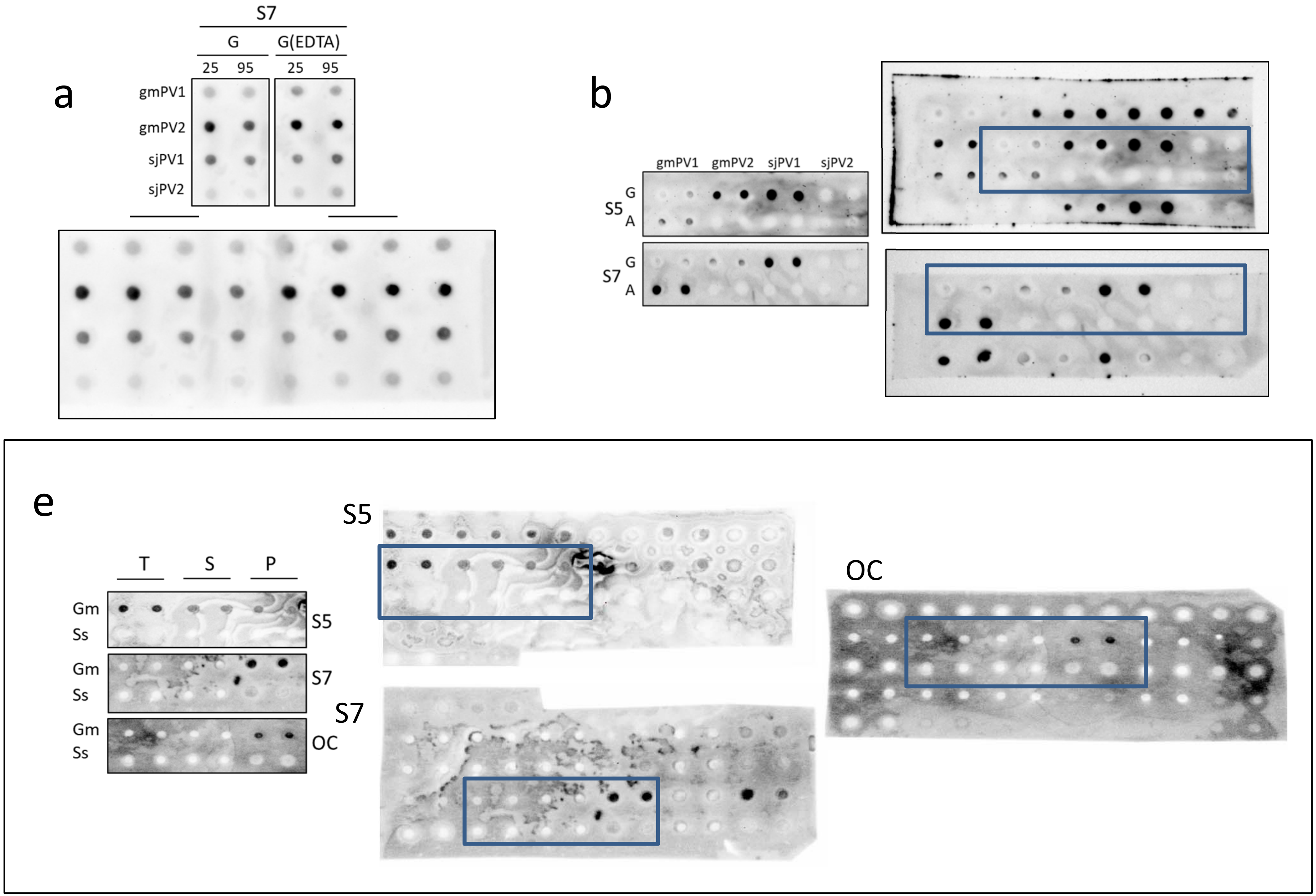
